# Revealing invisible cell phenotypes with conditional generative modeling

**DOI:** 10.1101/2022.06.16.496413

**Authors:** Alexis Lamiable, Tiphaine Champetier, Francesco Leonardi, Ethan Cohen, Peter Sommer, David Hardy, Nicolas Argy, Achille Massougbodji, Elaine Del Nery, Gilles Cottrell, Yong-Jun Kwon, Auguste Genovesio

**Author notes:** correspondence should be addressed to AG and YJK. equal co-contribution (Alexis Lamiable, Tiphaine Champetier).

## Abstract

Biological sciences, drug discovery and medicine rely heavily on cell phenotype perturbation and observation. Aside from dramatic events such as cell division or cell death, most cell phenotypic changes that keep cells alive are subtle and thus hidden from us by natural cell variability: two cells in the same condition already look different. While we show that deep learning models can leverage invisible features from microscopy images, to discriminate between close conditions, these features can yet hardly be observed and therefore interpreted. In this work, we show that conditional generative models can be used to transform an image of cells from any one condition to another, thus canceling cell variability. We visually and quantitatively validate that the principle of synthetic cell perturbation works on discernible cases such as high concentration drug treatments, nuclear translocation and golgi apparatus assays. We then illustrate its effectiveness in displaying otherwise invisible cell phenotypes triggered by blood cells under parasite infection, the presence of a disease-causing pathological mutation in differentiated neurons derived from iPSCs or low concentration drug treatments. The proposed approach, easy to use and robust, opens the door to the accessible discovery of biological and disease biomarkers.

## Introduction

Assessing quantitative differences in visual cell phenotypes from microscopy images finds many applications in biological basic research, drug discovery and medicine (Lundervold and Lundervold 2019; Moen et al. 2019; Ching et al. 2018). This task has historically relied on hand-crafted image analysis algorithms when the phenotype of interest was known and visible, and the cells could be segmented (van der Walt et al. 2014; Carpenter et al. 2006). Likewise, when the cells can be segmented but the phenotype is too complex for a discriminative function to be written, the latter can still be learned by training machine learning on vectors of handcrafted features, or directly from segmented cell images using supervised deep learning (Jones et al. 2009; Dürr and Sick 2016). Furthermore, when phenotypes are too subtle to be discernible, thus preventing supervised training, quantitative features can still be computed or learnt in a self-supervised fashion from segmented cells. These profiles of quantitative features can then be used to achieve complex tasks such as predicting the mechanism of action or the target of drugs (Ljosa et al. 2013; Rose et al. 2017). Ultimately, when the phenotype is too subtle to be discernible and the cells cannot be individually segmented -as for instance when dealing with cancer cell colonies, neurons, complex tissues made of fibroblasts, muscle cells or intra- and intercellular structures with unclear boundaries - a deep network model can often still be trained in a supervised fashion from raw image tiles, in order to discriminate conditions with good accuracy, bypassing the cell detection step. In other words, differences and similarities between cell phenotypes not discernible by the human eye can still, in many cases, be assessed by deep learning, demonstrating that microscopy images contain much more information than what can be seen (Scheeder, Heigwer, and Boutros 2018; Michael Ando, McLean, and Berndl 2017). It is a great opportunity for biology and medicine to be able to discover and investigate sounder, more subtle and broader classes of phenotypes and biomarkers. However, as the phenotype cannot actually be seen, these last approches offer no guarantee that the discrimination is not based on a bias, an experimental artefact or a biological phenotype irrelevant to the considered assay.

A solution to this issue consisted in attempting to explain *a posteriori* what such a classifier had learnt. This has historically been addressed by inference from the loadings of latent variables in the case of linear analysis, or from the decision rules in classification tree (Child 2006; Ziegel 2003). However, deciphering decision rules from most non linear classifiers, such as RBF Support Vector Machines or Neural Networks, has been notoriously difficult. With the advent of the deep learning era, the issue has worsened because the ability of deep networks to solve more complex problems came with a drastic increase in the number of parameters, due to the deep transformation needed to reach a representation relevant to the objective: the model works as a black box that performs the task well, but we lose track of how it does so (Xie et al. 2020). This issue has recently gathered significant attention, to the point where the very use of deep learning for high stakes decisions is currently being questioned (Rudin 2019). Furthermore, what would be a good description of how a model classifies is neither clear nor does it reach consensus across data types or domains. This problem is currently under debate and it seems that a good explanation should be a tradeoff between interpretability and completeness (Gilpin et al. 2018).

In the case of image data, compelling visualization methods to explain deep classifiers have been developed over the last 10 years. They can be roughly split into perturbation approaches that aim to infer important features by modifying the image input, and back propagation approaches that aim to specify which part of the input is related to a specific class. Perturbation methods comprise works such as occlusion sensitivity (Matthew D. Zeiler and Fergus 2014), representation erasure (Li, Monroe, and Jurafsky 2016), meaningful perturbation (Fong and Vedaldi 2017) and prediction difference analysis (Zintgraf et al. 2017). Back propagation methods comprise work such as activation maximization (Erhan et al. 2009), deconvolution (M. D. Zeiler et al. 2010), Layer-Wise Relevance Propagation (Bach et al. 2015), class activation mapping (Zhou et al. 2016), feature importance (Shrikumar, Greenside, and Kundaje 2017) and Integrated Gradients (Sundararajan, Taly, and Yan 2017). However, these works make the general assumption that visual feature input related to class output is visible, and therefore mainly needs to be pointed out in order to infer a meaning. Indeed, datasets are often taken from public repositories of natural image such as ImageNet where objects to be classified are clearly identifiable by eye and are even annotated this way (Deng et al. 2009). In the case of subtle phenotypic changes, this assumption is not met. In short, it is not sufficient to localize an area where discriminative features could stand in the image because, due to cell variability, these discriminative features can still not be seen.

In this work, instead of first training a classifier, and then attempting to explain what it has learnt, we propose a straightforward method based on established work that can directly provide a general description of the differences between any two or more image sets. This permits us to infer and explain invisible differences between conditions, without relying on intermediary classifier training.

Our approach was based on the hypothesis that if two cellular visual phenotypes can be distinguished by a deep network, but not by a human eye, this is mostly due to the fact that the cell to cell variability within an image largely overlaps the cell to cell variability between phenotypes, hence the former hides the latter. In short, natural variability prevents us from seeing the subtle phenotype difference between two images of cells. We then extrapolated that the subtle differences between two close phenotypes could be made visible and inferred if the natural variability was canceled out. To this end, instead of comparing two real images of cells under condition A and B that display necessarily different cells, we used a generative model to translate an image of cells under condition A to the image of the same cells under condition B. The signal modification thus produced could be defined as the difference between phenotypes induced by A and B. To implement this idea, we used conditional Generative Adversarial Networks (GANs).

GANs produced a drastic improvement in the image synthesis field by taking advantage of an adversarial training scheme taking place between a generative network and a discriminative network (Goodfellow, Pouget-Abadie, and Mirza 2014). This principle has since been widely used and improved in many ways, in order to generate various kinds of data. Numerous compelling works in image generation and translation have been proposed, including recent work explaining black box classifiers, but, to our knowledge, never with the aim of explaining invisible changes between conditions (Choi et al. 2018; Baek et al. 2020; Zhu et al. 2017; Lang et al. 2021; Singla et al. 2021). On the contrary, because the aim is different, domains chosen for such work were usually visually different in order to validate the approaches.

We thus show that conditional image synthesis enabled us to artificially reproduce the effects of various perturbations on a single cell image. We then validated on two standard visual assays that quantitative differences between real conditions could be retrieved from synthesized ones. Finally, we illustrated the possibilities offered by this approach for deciphering subtle variations of phenotypes triggered by a parasite infection on blood cells, a mutation in human neurons, or a low concentration of compound treatment, all invisible to the human eye.

## Results

### Conditional GAN can synthesize cell phenotype perturbations

We trained a conditional GAN (see Online methods) to reproduce the transformations - or translations - that cells undergo when subject to various high dosage compound treatments. In this way, using a single model and a single training, we could artificially produce images of the effect of various compound treatments, when applied to a single image of untreated cells. The real images of **Figure 1**A,B were taken from an assay of Human MCF-7 breast cancer cells treated for 24h with small molecules at various concentrations. After fixation, cells were labeled for F-actin, B-tubulin and DNA, and imaged by fluorescent microscopy, as described in (Caie et al. 2010). In this first training, images of treated cells were chosen so as to display an obvious phenotype at high concentration (**Sup. fig. 1**). Results demonstrated the capability of conditional image generation to reproduce cell phenotypes induced by compound treatment (**Figure 1**A). Furthermore, synthesis could be performed for all treatments from the same cell image, and latent traversals enabled us to see gradual changes that need to be performed in order to transform an untreated (DMSO) cell to any compound treated cell (**Figure 1**B). We engineered a web interface to ease visualization and manipulation of such data on many examples for all datasets considered in this work (https://www.phenexplain.bio.ens.psl.eu/). Interestingly, beyond the phenotypes themselves, the synthetically treated cell images displayed a lower cell count which is consistent with the fact that these treatments are toxic at high concentration (**Figure 1**A,B). Notably, as intermediate generated images are also validated by the conditional GAN discriminator as possible images from the dataset, cells on the border do not gradually disappear, but tend to gently shift out of the field of view when possible. This last example demonstrates that this type of morphing doesn’t match any sort of real dynamic. However, it does certainly help to visualize the differences between phenotypes, especially when they are subtle.

**Figure 1:**
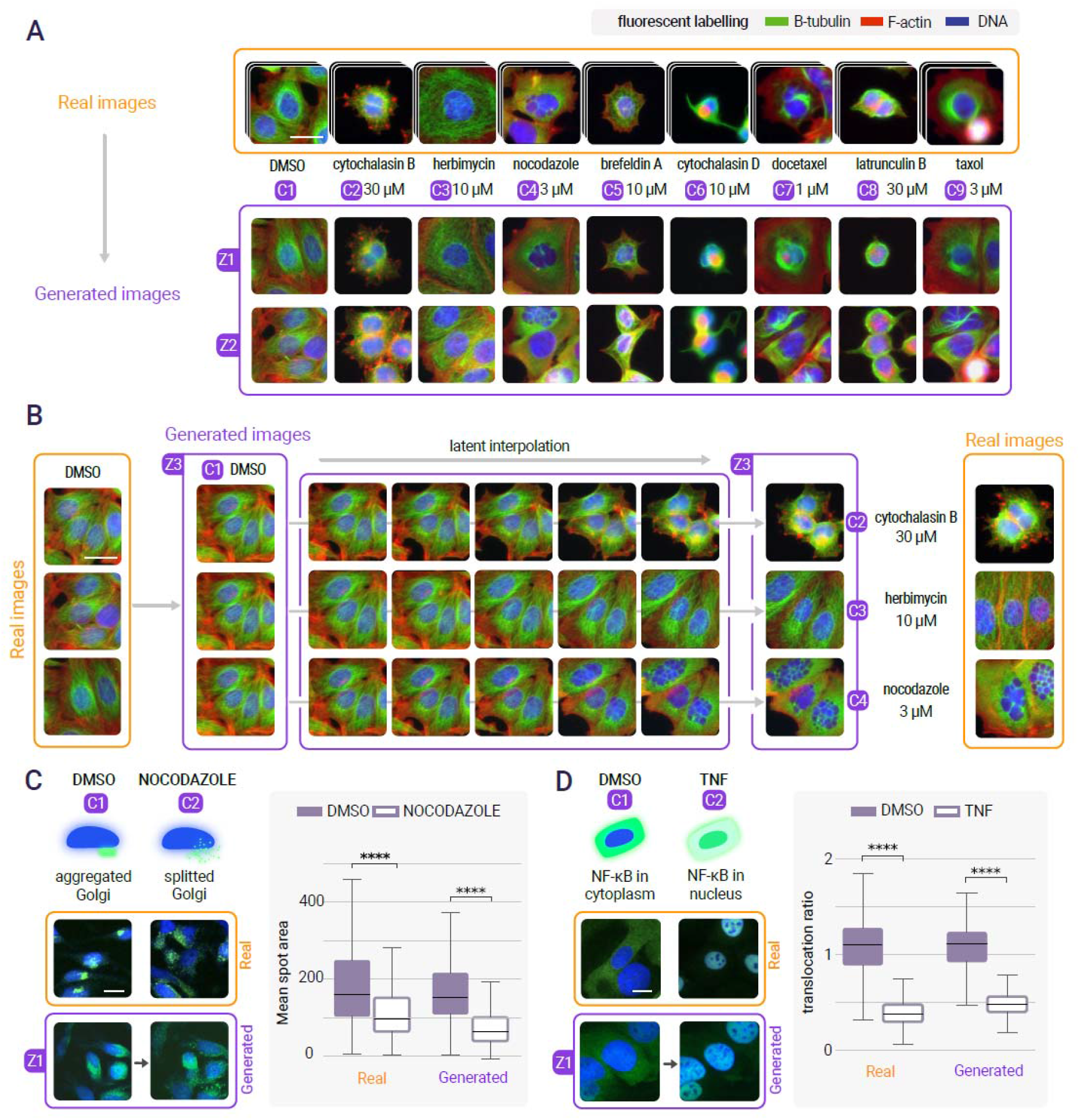
Conditional synthesis of cell phenotype perturbations. **A -** A conditional GAN is trained on real images of DMSO and high concentration drug treatments (C1, C2, C3, etc.), scale bar is 20_μ_m. It is then possible to generate synthetic images of any of these treatments from the same random seed, here z1 or z2. Synthetic images (violet) are indistinguishable from real ones (orange). As the compound concentrations are high on these exemples, the phenotypic changes are obvious and visible. Importantly, some surrounding cells in the negative control (DMSO) are removed from the images of compound treatments because of their toxicity. **B -** A latent traversal can then be computed for a single seed (z3) from the untreated state (C1=DMSO) to 3 different treatment effects (C2,C3,C4) displaying each a different gradual change of the same cells. **C -** A standard assay such as the nocodazole induced golgi scattering (green) can be reproduced with synthetic images, scale bar is 20_μ_m. An image analysis measurement (mean spot area) performed on real and synthetic images of both conditions led to the same quantitative conclusion (N=1000 for each distribution, two sided t-test). D - Another standard assay displaying TNF-induced NFkB translocation (green) can also be reproduced (N=1000 for each sampled condition, two sided t-test), scale bar is 20_μ_m.

### Synthetic cell features match those of real treatment

To quantitatively evaluate the ability of conditional image generation to properly reproduce real phenotypes, we applied it on two assays in which phenotypic alteration could be independently quantified using standard image analysis tools. The first assay monitors the morphology of the golgi apparatus state. By treating cells with nocodazole, microtubules are depolymerized and the golgi originally located at about the center of the cell is split into mini stacks (**Sup. fig. 2**). A simple average spot size difference measured on 1000 real treated and 1000 untreated images could be reproduced when computed only from synthetic images (**Figure 1**c). The second assay monitored the subcellular location of NF-kB (nuclear factor kappa B) protein. When treated with TNFα (pro-inflammatory cytokine tumor necrosis factor alpha) the transcription factor translocates to the nucleus, and the corresponding fluorescence signal moves from the cytoplasm to the nuclear area, with cells displaying bright green nuclei (**Sup. fig. 3**). Similarly, for this assay, the difference of nucleus versus cytoplasmic fluorescence ratio per cell measured on 1000 real treated and 1000 untreated images could be reproduced very closely when computed from synthetic images (**Figure 1**d). Confirmation that quantitative features difference on real images could be reproduced on synthetic images for both assays suggested that imperceptible yet reliable phenotype variations could also be reproduced.

### Deciphering blood cell changes triggered by a parasite infection

The gold standard diagnosis method performed on patients with suspected malaria in endemic areas is based on microscopic observation of blood film to detect intraerythrocytic *plasmodium spp*. This method presents the compelling advantages of being inexpensive, rapid, not requiring any advanced device and usable directly on the field, close to the patients. However, this approach is also less sensitive than molecular ones such as qPCR that is more expensive, necessitates lab work and an expensive device, and is not available on the field, however it can reveal “submicroscopic active infection”. This is defined as a qPCR positive result while no visible parasites were found by microscopic examination and can represent up to half the positive cases in some areas (data not shown). We then aimed to evaluate the hypothesis that such infection could still be detected from microscopy images despite the absence of visible parasites. This would pave the way to a possible fast and low cost microscopy-based sensitive diagnostic test available on the field. We then collected thin blood smears and selected on one hand 100 slides that were diagnosed positive for plasmodium infection by qPCR but did not display any visible parasites, thus classified as negative by a microscopist, and on the other hand 100 slides that were diagnosed negative by qPCR, therefore non-infected patients (**Figure 2**A). 300 images were extracted from each of these 200 slides and visual inspection confirmed that no parasite nor visible features could distinguish the qPCR+ from the qPCR-cases (**Sup. fig. 4**). However, training a simple convolutional neural network showed that classification of these slides could be achieved with 74% accuracy (**Sup. fig. 8**), indicating that it contained invisible discriminative features. We then trained a conditional GAN on these close phenotypes. Results show that transforming an image of qPCR-to qPCR+ necessitates generation of more anemia cells. Indeed the translation from a blood cell of a healthy patient to the blood cell of an infected patient seems to wipe a part of the hemoglobin content which, while not specific to the malaria infection, tends to denote an infectious context (White 2018; Scovino, Totino, and Morrot 2022). Furthermore, we also found that some cells were transformed into crenated cells which were previously reported to be related to certain liver diseases (Privitera and Meli 2016). Finally, the system also detected that there might have been a relative excess of color staining in some of the negative slides because the synthetic translations to the positive case showed a lower amount of debris due to the staining step (**Figure 2**B).

**Figure 2:**
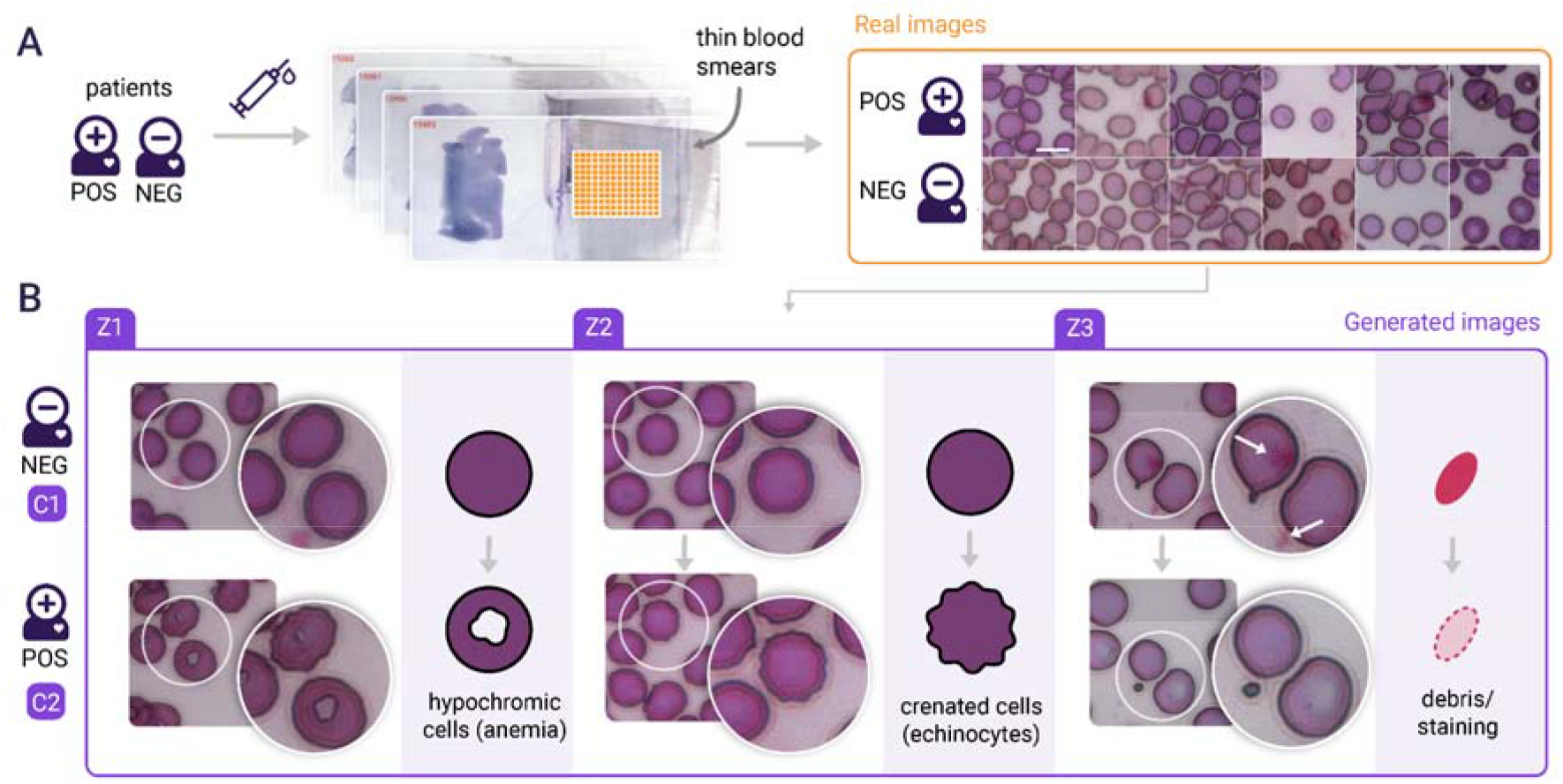
Unraveling red blood cell morphological changes related to an infectious environment. **A** - Images of blood cells were extracted from thin blood smears sampled from a population of people exposed to malaria. 200 slides were selected as negative to Malaria by microscopists, meaning that no parasites could be found on any of these slides, with nevertheless half of them found positive by qPCR. Note that on both qPCR positive and qPCR negative slides, images extracted displayed variable cell densities with variable background, did not contain any parasites, and did not show any identifiable systematic visible differences, scale bar is 10_μ_m. **B** - In order to identify discriminative features between qPCR+ and qPCR-slides, we used 60,000 such images from these 200 slides to train a conditional GAN. This panel shows three representative generated images of the results found. Z1 displays a visual difference that can be interpreted as an increase of anemia: the content of some blood cells lose hemoglobin (displayed as a hole or a white halo in the cell). This phenotype could barely be identified from real images data because both qPCR+ and qPCR-slides contain anemia cells. Additionally, Z2 displays some deformations of the cell membrane producing crenated cells. Finally, Z3 shows that the negative sample contained more debris due to staining as a translation to qPCR+ tends to remove these artifacts. Indeed, the system cannot discriminate between relevant differences of phenotypes from biologically irrelevant differences related to possible technical or experimental biases.

### Uncovering morphological variations in patient derived dopaminergic neurons

Subsequently, we wondered if this approach could be used to investigate subtle morphological variation induced by a mutation. We used iPSCs reprogrammed from fibroblasts of a Parkinson’s disease (PD) patient carrying the LRRK2 G2019S mutation along with an isogenic control, in which the mutation was genetically rescued to the wild-type sequence by gene editing and differentiated the iPSCs into dopaminergic neurons (Vuidel et al. 2022). The LRRK2 G2019S mutation, localized in exon 41 of the LRRK2 (leucine-rich repeat kinase 2) gene, is causally linked to the development of Parkinson’s Disease (Tolosa et al. 2020). After immunostaining for nuclei, Tyrosine Hydroxylase (TH, a marker for dopaminergic neurons) and alpha synuclein (SNCA, a protein that accumulates in Lewy bodies and Lewy neurites in Parkinson’s disease and other synucleinopathies), we acquired images of isogenic dopaminergic neurons with and without the G2019S mutation. Images of dopaminergic neurons obtained in these two conditions looked indistinguishable and could not be segmented with image analysis tools (**Figure 3**A and **Sup. fig. 5**). However a convolutional network could discriminate them with 63% accuracy at the single image level and with 100% accuracy when aggregated at the well level (**Sup. fig. 8**). Interestingly, training a conditional GAN on these two image sets highlighted differences between conditions from which interpretation was possible (**Figure 3**B). The generated images of engineered WT cell cultures displayed an increase of dopaminergic neurons, a decrease of non-differentiated cells and an increase of alpha synuclein in some of the remaining non-differentiated cells as compared to the LRRK2 G2019S cultures. These differences denoted an increase of the differentiation efficiency in one of the conditions which could be due either to the suppressed mutation or to an experimental bias due to the genome edition for instance.

**Figure 3:**
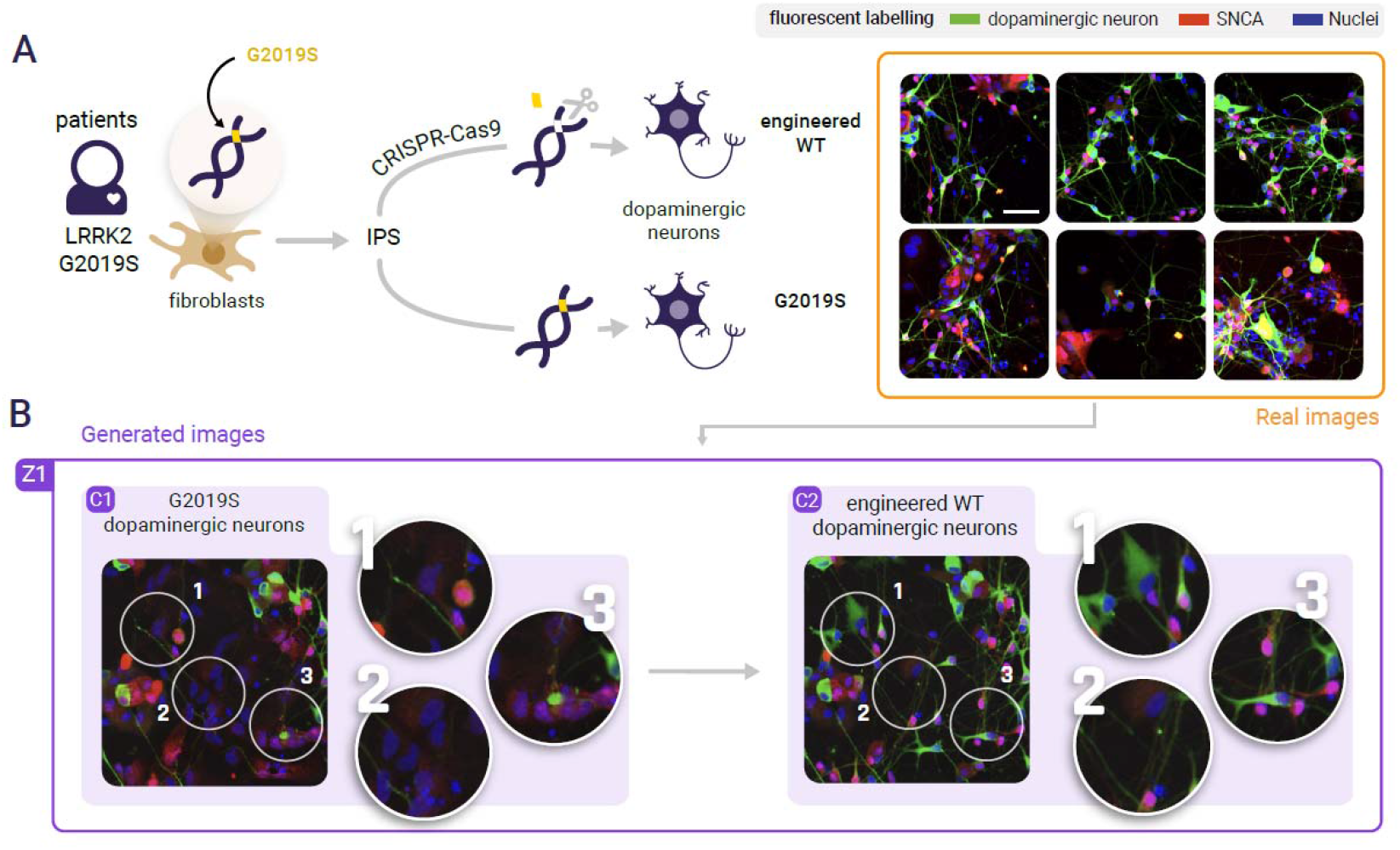
Unraveling invisible morphological variation in a patient derived dopaminergic neuron assay. **A -** IPSCs are derived from fibroblasts sampled from a LRRK2 G2019S mutated Parkinson’s patient. These IPScs are then reprogrammed to dopaminergic neurons with or without a CRISPR-cas9 correction of the G2019S mutation. The last is then an isogenic engineered wild type. Large sets of confocal images with one dye labeling for Nuclei, and antibody stains labeling for alpha synuclein and TH cells, scale bar is 20_μ_m. Real images display no detectable visual systematic differences. **B -** A conditional GAN trained in order to identify differences between these two close conditions displays 1 - an increase of dopaminergic neurons in the engineered WT condition: part of the IPScs translate into neurons, 2 -removal of some of the IPScs, 3-accumulated alpha synuclein in the nuclei of some of the remaining IPScs.

### Low drug concentration effects can be made visible

We then used the same approach to display the invisible morphological changes that could be induced by low concentration drug treatment. To this end, we considered images with cells treated at very low concentration of the same drugs used for **Figure 1**. Due to cell variability, the real images of these treatments displayed no visible differences both compared to the untreated cells and between one another (**Sup. fig. 6**). However, for most of these treatments, a simple convolutional network could be trained to distinguish both cases (**Sup. fig. 8**). We then trained a conditional GAN to translate images of untreated (DMSO) cells to images with cells treated with low concentration of these drugs (**Figure 4**A). Interestingly, results showed that any of these treatments, even at the lowest available concentration, had a slight toxic effect, as a few cells on the border of the images were systematically removed. Furthermore, some treatments, such as cytochalasin D, seemed to systematically contract cytoplasm, while others, such as taxol or nocodazole, extended it (additional examples can be manipulated with our web interface).

**Figure 4:**
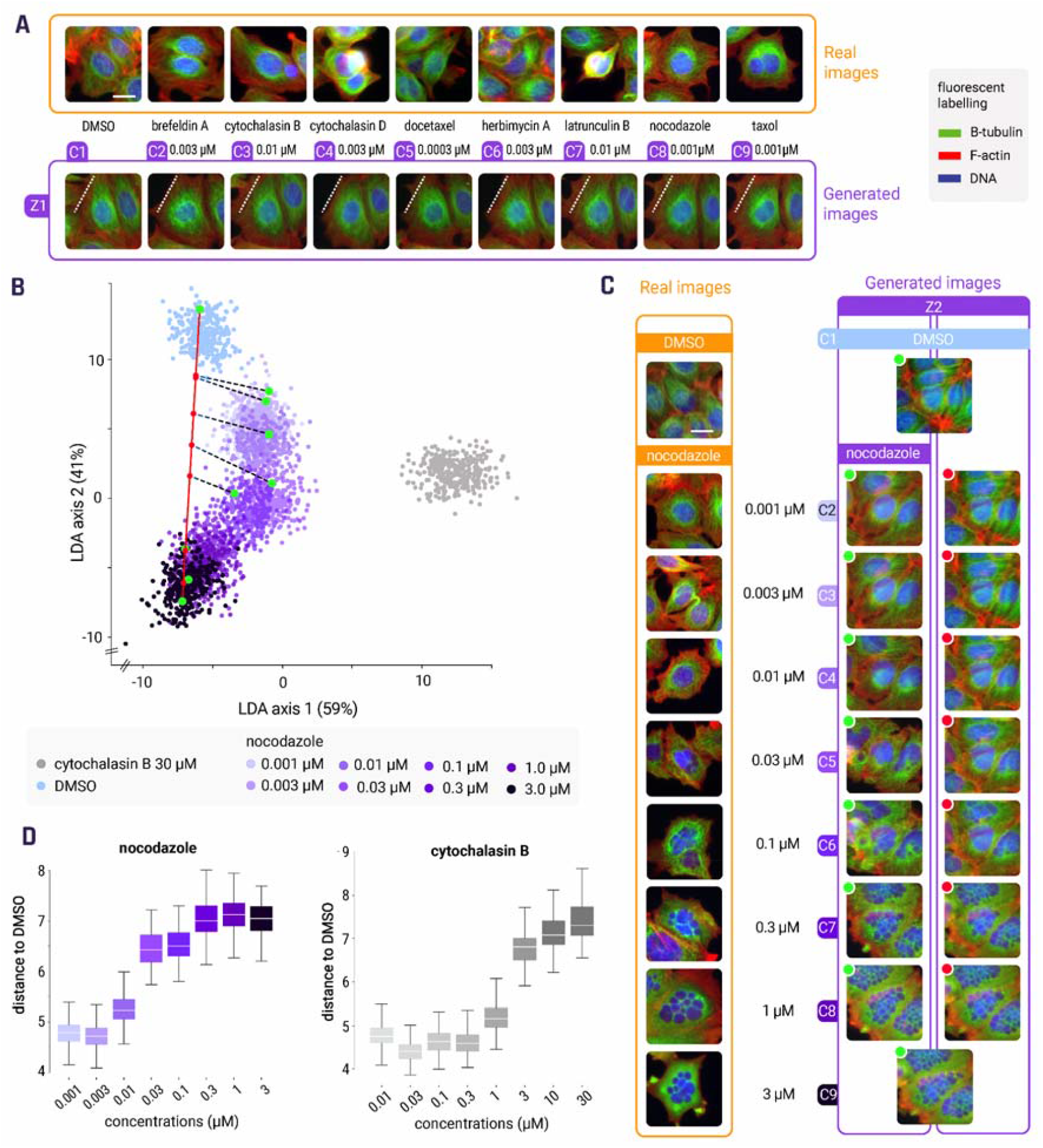
Morphological effect of perturbations by low dose compound treatments and dose response. **A -** The same compounds used at high concentration in Figure 1 were also plated at very low concentrations. The corresponding images cannot be visually distinguished from untreated cells (DMSO) and from one another (first row), scale bar is 20_μ_m. A conditional GAN was trained on real images of these low concentration compound treatments. By doing so we could generate artificial images of these perturbations on the same cells and compare them with DMSO and between each other (second row). We see that most treatments, even at low doses, had a slight toxic effect as they removed cells on the image borders compared to DMSO. Furthermore, some compounds tend to expand the cell cytoplasm while some others contract it. **B -** An LDA plan is computed from generation of 3000 DMSO, 3000 Nocodazole and 3000 cytochalasin B at highest doses. Then, 300 samples of each available concentration of the Nocodazole treatment were drawn and projected onto this plan. **C** - Left column, real images from a nocodazole dose response, middle column, z2 is a random seed used to generate perturbations of the same cells for each concentration (green dots on panel B) and, right column, z3 represent their orthogonal projections on the latent traversal between DMSO and the highest nocodazole dose in the W space: red dots on the red axis. **D** - Computation of the distances in the W space of 1000 samples from each dose (C2, C3, etc.) to the DMSO (C1) for a given compound enables construction of a dose response curve describing the gradual intensity of the morphological changes.

### Dose response can be approximated by latent traversal

While the sequence of images generated from the latent interpolation between two conditions could not be considered as matching any dynamic reality, we wondered whether it could approximate treatments at variable concentrations, or a so-called dose response. To evaluate this, we computed a Linear Discriminant Analysis (LDA) from 3000 data points generated for each of 3 extreme conditions: nocodazole and cytochalasin B at highest concentrations and DMSO. We then projected 300 data points from all concentrations of Nocodazole on this discriminative plan. The result shows that the consecutive concentrations do form a clear path on the discriminative plan from DMSO to the highest dose (**Figure 4**B). We could then, for a given seed z, display side by side the generated sequence conditional to each dose on one hand, and their orthogonal projection (in the high dimensional space W) on the latent traversal on the other hand (**Figure 4**C). The images thus retrieved for several compounds suggested that the latent linear interpolation between DMSO and a high dose was a reasonable approximation of the dose response. We then measured the distance in the latent space between the untreated cells (DMSO) and each generated dose of nocodazole and cytochalasin B (**Figure 4**D). Interestingly, the lowest doses could already be discriminated from DMSO, and the distance to DMSO increased with the concentration of compound, suggesting a method to quantify the relative amplitude of the effect of compound treatments on a considered assay, without performing any specific measurement.

## Discussion

Assessing cellular state alteration is paramount to cell biology. Undoubtedly, understanding the subtle effect of genetic, chemical or disease-based perturbation on cell phenotypes is instrumental. While deep learning offers compelling tools to discriminate between two imaged conditions, even when no differences are perceptible, it is hard to figure out if differences between two image sets are related to a relevant phenotypic variation.

In this work, we suggest that cell variability is the main barrier to visually assessing the subtle effects produced by a perturbation on cells or simply the difference between two close conditions or mutants. We thus proposed an approach to make the translation of cell images between two close phenotypic conditions possible and quantitatively assessed that the difference obtained corresponds to the actual modification. We then demonstrated the advantage of such a method in deciphering fine phenotypic alteration triggered by an infection, a mutation on patient-derived neurons or low concentration of drugs, otherwise invisible. We further showed that the latent space thus created enabled quantitative comparison between close imperceptibly different perturbations, such as low doses of a compound treatment, and permitted the approximation of a dose response.

Interesting solutions such as class activation maps do exist to explain what a black box classifier has learnt. They essentially point to areas where visible discriminative features lay on an image (Zhou et al. 2016). However, when these features are invisible, such as for subtle cell phenotypes, these methods become inefficient and cannot lead to interpretation, as shown on our own data (**Sup. fig. 9**). They could not lead us to use real images to even partially reach the clear observation we could make otherwise from synthetic ones. Some recent work has used generative models to explain what a classifier learnt still by pointing out already perceptible or known features (that were actually used to annotate the pictures), but, to our knowledge, never evaluated on invisible cell phenotypes in the context of various assays (Lang et al. 2021; Singla et al. 2021). Furthermore, our approach does not focus on explaining a trained classifier and therefore does not require one. On the contrary, these methods attempt to explain what a classifier can discriminate against, and not directly what the differences between datasets are. While these goals are different, using a trained classifier toward our end goal would add a non neutral intermediate proxy to assess the difference between phenotypes. Indeed, any hyper parameter of the classifier, such as its capacity or even the way it is trained, would add its form of bias between the images and our actual understanding of what the difference between conditions might be. On the contrary, our approaches offer to highlight the signal difference that discriminates between two close phenotypes directly, without the need for an intermediate classifier.

What could be seen as a current limitation of conditional GAN to explain the variation between invisible phenotypes is that it cannot be used to artificially alterate a real image. Indeed, instead of using a conditional GAN, an unconditional GAN along with GAN inversion could be used to obtain phenotype translation (see method 2 in online methods). However, attempting to use several GAN inversion methods to retrieve latent code from a real image in a GAN or in a conditional GAN latent representation led to unsatisfactory image results on visible cell phenotypes (Richardson et al. 2021; Alaluf, Patashnik, and Cohen-Or 2021). Concretely, even some high concentration perturbation images could not be reconstructed properly by swapping important details and organel positioning, which we considered problematic in identifying subtle phenotypes. Many reasons can explain these results as assessed in a recent review on GAN inversion which is still an open problem (Xia et al. 2021). On the other hand, we demonstrated here that performing transformation on real images was definitely not required in order to decipher variations of a subtle phenotype.

Another apparent constraint of the proposed approach is that it still requires a human to infer the difference from the visual transformation and does not discriminate relevant from irrelevant differences on its own. While this could be seen as a limitation, it in fact indicates that the system is unbiased. Datasets do include differences and a conditional GAN will just display them without favoring relevant biological features over unwanted biases. Using such an approach, we could then reverse the classifier explanation paradigm, and rather than explaining a posteriori a classifier trained blindly, we could envision first extracting relevant and irrelevant differences between two conditions from a dataset using this approach, and then building an efficient guided system that would consider only the features of interests, disregarding the identified biases or irrelevant features. The staining artifacts in the red blood cell example, due to the fewer number of slides in the training set, is typically such an irrelevant feature that we would rather not want a classifier upon which to base its discrimination.

An extension of this work could consider disentangling the image transformation into interpretable factors of variation so that the weight of quantitative features may be more concretely established and used afterward.

This method is straightforward to use, can be applied to any microscopy image modalities and is accessible to the community. Beyond displaying invisible phenotypic variations, this system proposes a straightforward way to discover yet unknown biomarkers.

## Online methods

### Conditional GAN

A cell image transformation between phenotypes can be obtained using a conditional GAN by producing a latent traversal of a single random seed between two trained conditions. Any GAN architecture can be made conditional by modifying the regular GAN minimax game:

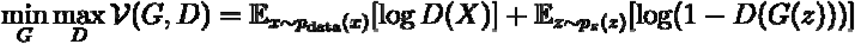

by introducing auxiliary information both in the generator G and the discriminator D:

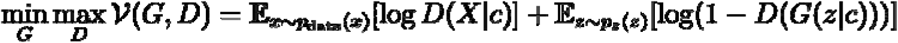

In practice, a linear layer is used in both the generator networks G and the discriminator D to embed each condition as a learnable vector. This embedding vector is then concatenated to an early layer of the generator. Similarly, a condition embedding vector is concatenated or multiplied to the output of one of the deepest layers of the discriminator. At training time, the conditional generator is trained to fool the conditional generator for each condition. The trained generator acts as a bank of unconditional GANs but presents the advantage of sharing the same latent representations.

### Translation and latent traversal

The output of each layer of a deep network is a latent representation of the network input. Any displacement or translation in one of these latent spaces in a conditional or unconditional GAN will produce an image transformation as output. We define the phenotype translation as the unique translation that transforms the same random seed z (a synthetic image of a cell) from a condition c_1_ to a condition c_2_. Also, when considering subtle phenotypes, we found it very useful and insightful in practice to actually compute and display the latent traversal, that is the consecutive images showing the transformation from one phenotype to the other. While the gradual transformation dynamic is not biologically relevant, each image is a possible intermediate state because it was assessed as realistic by the discriminator. Furthermore, it allows us to gradually see the modifications of fine features that need to occur in order to transform an image from one condition to another. This is especially interesting in the case of subtle phenotypes, when conditions are very close to each other.

We have identified three ways to generate latent traversal between conditions that can each be used for different purposes.

**The first approach** can be used with any conditional GAN architecture and simply consists of a linear interpolation between the embedding vectors of two conditions for the same random seed. Given a generator G that includes an embedding layer E that maps a condition c to a vector e and the remaining layers F that takes a random seed z and an embedding vector e, then G(z,c) = F(z,E(c)). A latent traversal of N+1 images can then be computed from the output of the embedding layer in this way:

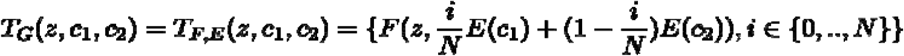

**The second approach** is specific to StyleGAN2, an efficient architecture used here that builds an intermediate latent representation named W that, by construction, tends to be disentangled. It is then possible to directly interpolate in the W space, which is interesting as it also allows us to compute latent traversals without prior annotations, which is useful if one wishes to modify real unannotated image after GAN inversion, compute distances to obtain dose responses curves, or perform projections, as it is done in the manuscript. StyleGAN2 is composed of a mapping network M and a synthesis network S. A random seed z and a condition c are combined to obtain the intermediate representation w=M(z,c), then an image can be synthesized with S(w). The centroid of the distribution of the intermediate representations of a treatment c_j_ can be estimated by simply computing the sample mean of K (e.g. 100,000) random seeds:

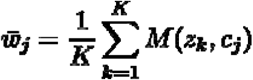

Applying the distribution shift from c_1_ to c_2_ to a given seed z can then be computed in this way:

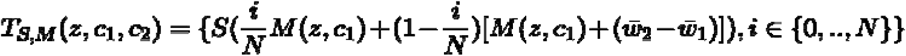

This approach can also be used with unconditional GANs to find axes of variations by inverting sets of images rather than synthesizing them.

**The third approach** is also specific to StyleGAN2 and can only work with generated images. A latent traversal corresponding to a phenotype translation can be obtained in a straightforward way by considering linear interpolation between the locations of the same random seed z conditioned by c1 and c2 in W:

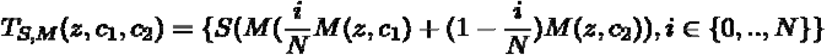

Note that these 3 approaches led to very close results; we thus mostly used the third approach in this work since for the sake of explanation we only need to manipulate synthetic images, and we did choose to use StyleGAN2 which is currently considered one of the best GAN by the computer science community.

### Phenexplain web interface

For ease of understanding and visualization, we created an interactive web interface with data produced from all the datasets we used in this work, making it possible to explore and manipulate more examples than the one presented in the figures (https://www.phenexplain.bio.ens.psl.eu/). For all the datasets, latent traversal can be explored interactively for a selection of examples. For the BBBC021 dataset, which provides more conditions, a more detailed exploration is possible. First, a linear discriminant analysis (LDA) was used to provide a projection of the latent space that properly separates three conditions (DMSO, and nocodazole or cytochalasin B at a high concentration). Their respective regions in the latent space can be explored to show that a variety of images can be generated for each condition. Latent traversal from DMSO to a compound can be performed either through a linear interpolation between DMSO and the highest concentration, or by successive interpolations between intermediate concentrations. We expect the latter to provide a trajectory that is more accurate, but the projection of these points onto the linear interpolation shows that the former is a reasonable approximation.

### Golgi assay

HeLa cells stably expressing EGFP-CCR5 were kindly provided by F. Perez’s team, UMR144, Institut Curie) and cultured as described in Boncompain et al., 2019 (Boncompain et al. 2019). Briefly, cells were grown in Dulbecco’s modified Eagle’s medium (DMEM) (Thermo Fisher Scientific) supplemented with 10% fetal calf serum (FCS), 1 mM sodium pyruvate, and penicillin and streptomycin (100 μg/ml) (Thermo Fisher Scientific). For the Golgi assay, 5.0 × 10^3^ cells were seeded on black clear-bottom 384-well plates (ViewPlate-384 Black, PerkinElmer) in 40 μl of complete medium. Twenty-four hours after cell seeding, DMSO (control solvent) and nocodazole were transferred robotically to plates containing cells to a final concentration of 10 μM and 0.5% of DMSO. After 90 min of incubation, cells were treated with 40 μM biotin for 120 min at 37°C. Cells were processed immediately after biotin treatment for immunofluorescence. Briefly, cells were fixed with 3% paraformaldehyde for 15 min and quenched with 50 mM NH_4_Cl in phosphate-buffered saline (PBS) solution for 10 min. Nuclei were counterstained with DAPI (Life Technologies) for 45 min.

### NF-κ B nuclear translocation assay

HCC1143 cancer cells (ATCC) were cultured with RPMI 1640 medium (Thermofisher) supplemented with 10% bovine fetal serum, 100□U/mL penicillin and 100□μg/mL streptomycin in a humidified environment consisting of 95% air and 5% CO_2_ at 37°C. Cells were seeded on black clear-bottom 384-well plates (ViewPlate-384 Black, PerkinElmer) in 40 μl of complete medium at density of 1000 cells/well for 24 h before exposure to 20 ng/ml TNFα for 30 min. After the incubation period, cells were fixed with 4% (vol/vol) buffered paraformaldehyde solution (PFA, Sigma-Aldrich) for 15 min, quenched with 50 nM NH_4_Cl solution, and permeabilized with 0.5% (vol/vol) Triton X-100 in PBS for 5 min. Cells were stained with Rabbit anti-human NF-κB p65 (C-20) (Santa Cruz Biotechnology) for 1 h and anti-rabbit A488 secondary antibodies together with 4’,6-diamidino-2-phenylindole dilactate (DAPI) for labeling nuclei. For both assays, image acquisition was performed using an INCell 2200 automated high-content screening fluorescence microscope (GE Healthcare) at a ×20 magnification (Nikon 20×/0.45). Four randomly selected image fields were acquired per wavelength, well, and replicate experiment.

### Isogenic iPSC lines

The iPSC lines used in this paper were generated by a third party and are described in detail in (Lee et al. 2020). The iPSC lines are deposited in the European Bank for Induced Pluripotent Stem Cells (EBiSC, https://cells.ebisc.org/) and listed in the Human Pluripotent Stem Cell Registry (hPSCreg, https://hpscreg.eu/). The original generators have obtained the informed consent from the donors. iPSCs were cultivated on Geltrex-coated (Thermo Fisher Scientific) dishes in StemMACS iPS-Brew XF (Miltenyi Biotech). The medium was changed daily, and cells were passaged twice a week using 0.5 mM EDTA in PBS (Thermo Fisher Scientific). Mycoplasma testing was performed twice per month.

### Differentiation of dopaminergic neurons from iPSCs, immunohistochemistry and image acquisition

Dopaminergic neurons were differentiated from iPSCs using a modified protocol based on Kriks *et al*. which is described in more detail in Vuidel et al. (Kriks et al., 2011; Ryan et al., 2013; Weykopf et al., 2019; Vuidel et al., 2022). Differentiated dopaminergic neurons were cryopreserved at 30 days in vitro (DIV30). For further experimentation, cryopreserved neurons were thawed in a water bath and centrifuged (400g, 5 min, RT) in basal medium supplemented with ROCK inhibitor (Miltenyi, #130-095-563). Cell pellets were resuspended in differentiation medium supplemented with a ROCK inhibitor. Media compositions are detailed in Vuidel et al., 2022. 384-well plates (Perkin Elmer, #6007558) were coated with 15 μg/ml Poly-L-Ornithin for 1 hour at 37 °C followed by 10μg/ml Laminin overnight at 4 °C. Using Tryphan Blue (Sigma, # T8154-20ML) and a Countess automated cell counter (Invitrogen), 10,000 cells/well were seeded in 384-well plates. Edge wells were avoided for seeding and filled with PBS. Thawed cells were incubated at 37 °C and 5 % CO2 for seven days until 37 DIV with differentiation medium changes every other day. Cell fixation and immunohistochemistry were performed as detailed in Vuidel et al., 2022. All imaging experiments were performed on a Yokogawa CV7000 microscope in scanning confocal mode using a dual Nipkow disk. 384-well plates (Perkin Elmer, #6007558) were mounted on a motorized stage and images were acquired in a row-wise “zig-zag” fashion at RT.

### Malaria microscopic and qPCR diagnoses

Whole blood specimens sampled on EDTA were initially collected during a population survey of asymptomatic patients for malaria infection in Benin. From each blood sample, approximately 8μl and 2 μl of whole blood were carefully placed on a microscopic slide to respectively perform thick and thin blood film for microscopic examination. Thick blood film was stained by giemsa 8% after drying whereas thin blood film was stained with conventionnal may-grunwald-giemsa after fixation by methanol. Microscopic examination was performed by a skilled microscopist and was considered as negative if no circulating parasites had been observed on the entire thick blood film or after microscopic screening of 40000 red blood cells on the thin blood film. If parasites were detected by microscopy, parasite load was estimated in parasites per microliter considering respectively 8000 leukocytes or 46000 red blood cells per microliter on thick and thin blood film. For the purpose of this study, a portion of the thin blood film was scanned using the AxioScan Z1 (Carl Zeiss Meditec) system at 40X and retrieved 20,000 images of size 256×256 pixels per slide. 300 out of 20,000 images per slide were randomly selected for deep network training.

Molecular investigations were performed using RT-qPCR. Briefly, for each patient, parasite DNA was extracted from 200 μL of whole blood using DNA mini kit as recommended by the manufacturers (QIAGEN®) and eluted in 60 μL of buffer. Parasite DNA genomic was detected by a screening biplex RT-qPCR simultaneously targeting the 18S gene of *P. falciparum* and *Plasmodium* spp in one well and *P. ovale* and *P. malariae* 18S gene in a second well. PCR conditions were: denaturation 10 minutes at 95°C following by 40 cycles of amplification composed by denaturation at 95°C 5 s and hybridation 1 min at 60°C. *Plasmodium falciparum* parasite load (parasite/μL) was estimated using a standard calibration curve obtained with 3D7 *P. falciparum* strain DNA extracted from blood pellet with known parasite load.

## Code availability

The step by step notice to train and use a conditional GAN on image datasets of biological conditions and performing latent traversal is available as Python/PyTorch scripts from github (https://github.com/biocompibens/phenexplain/).

## Acknowledgements

This work has received support under the program «Investissements d’Avenir » launched by the French Government and implemented by the ANR, with the references: ANR-10-LABX-54 MEMO LIFE ANR-11-IDEX-0001-02 PSL* Research University, Inserm ITMO Cancer 2021 - PHENEXPLAIN. TC was co-funded by ANRT and Ksilink under the Cifre program. This work was granted access to the HPC resources of IDRIS under the allocation 2020-AD011011495 made by GENCI. We thank the computing service of IBENS, Nikita Menezes for her help in assembling the figures and Mary-Ann Letellier for editing the manuscript.

## Ethical aspect

The iPSC lines used in this paper were generated by a third party and are described in detail in (Lee et al. 2020). The iPSC lines are deposited in the European Bank for Induced Pluripotent Stem Cells (EBiSC, https://cells.ebisc.org/) and listed in the Human Pluripotent Stem Cell Registry (hPSCreg, https://hpscreg.eu/). The original generators have obtained the informed consent from the donors. The malaria slides come from a survey carried out in the field in Benin with participants who gave their informed consent to participate. The study received approval from the institutional ethics committee of the Center for Research in Entomology of Cotonou n°023/CREC/CEI-CREC/SA

## Competing interests

The authors declare no competing interests.

## Notes

https://www.phenexplain.bio.ens.psl.eu/hd.html

### Competing Interest Statement

The authors have declared no competing interest.

