## Supplementary figures for "Revealing invisible cell phenotypes with conditional generative modeling"

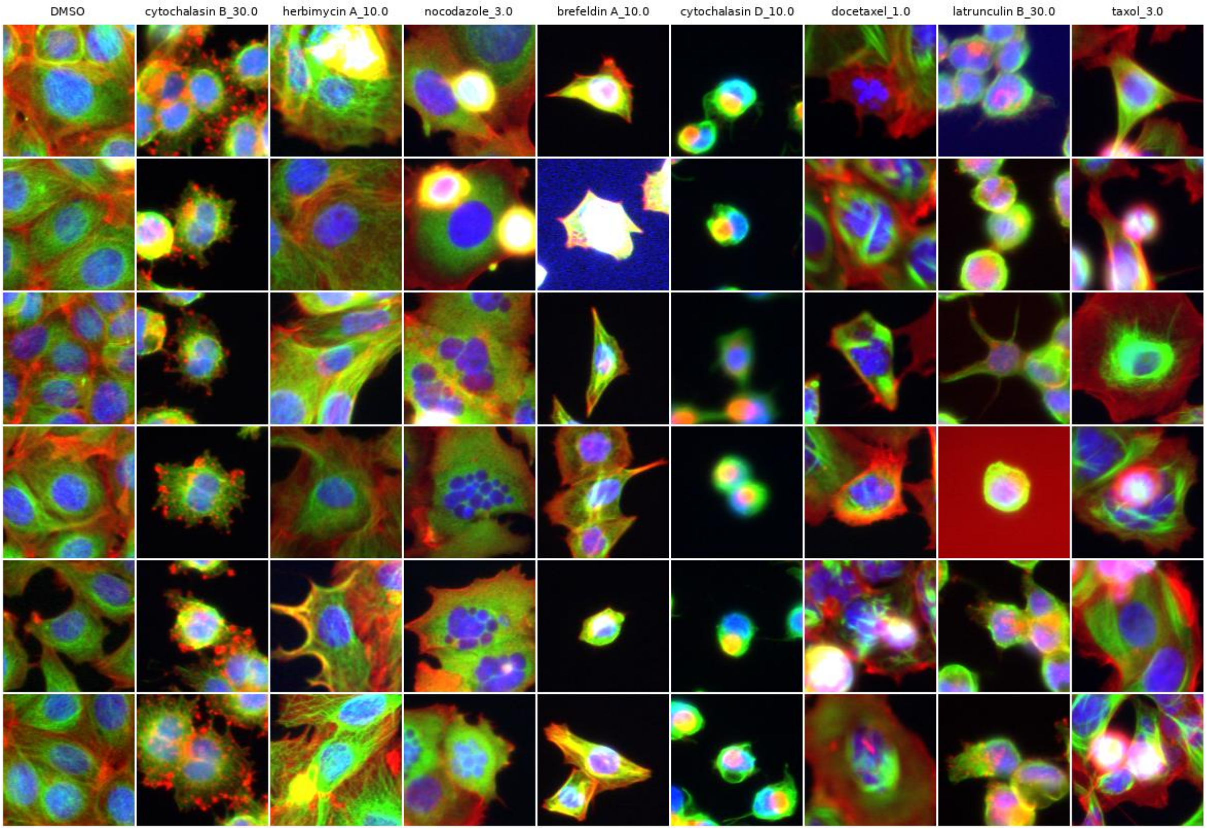


**Supplementary figure 1** Random samples of real cropped images of cells from high concentration compound treatments from the BBBC021 dataset (concentration unit is μM).

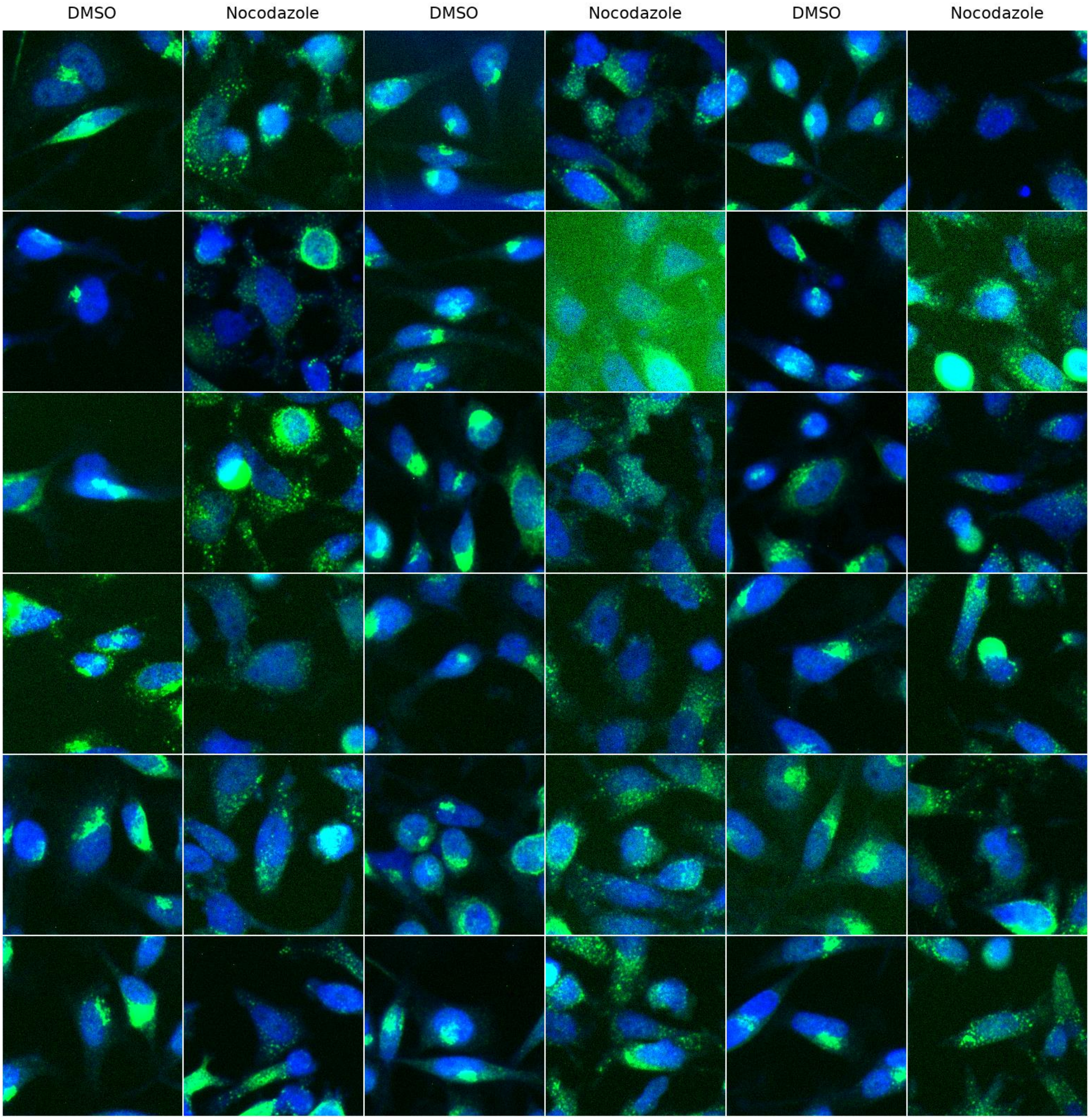


**Supplementary figure 2** Random samples of real cropped images of cells from the Golgi assay

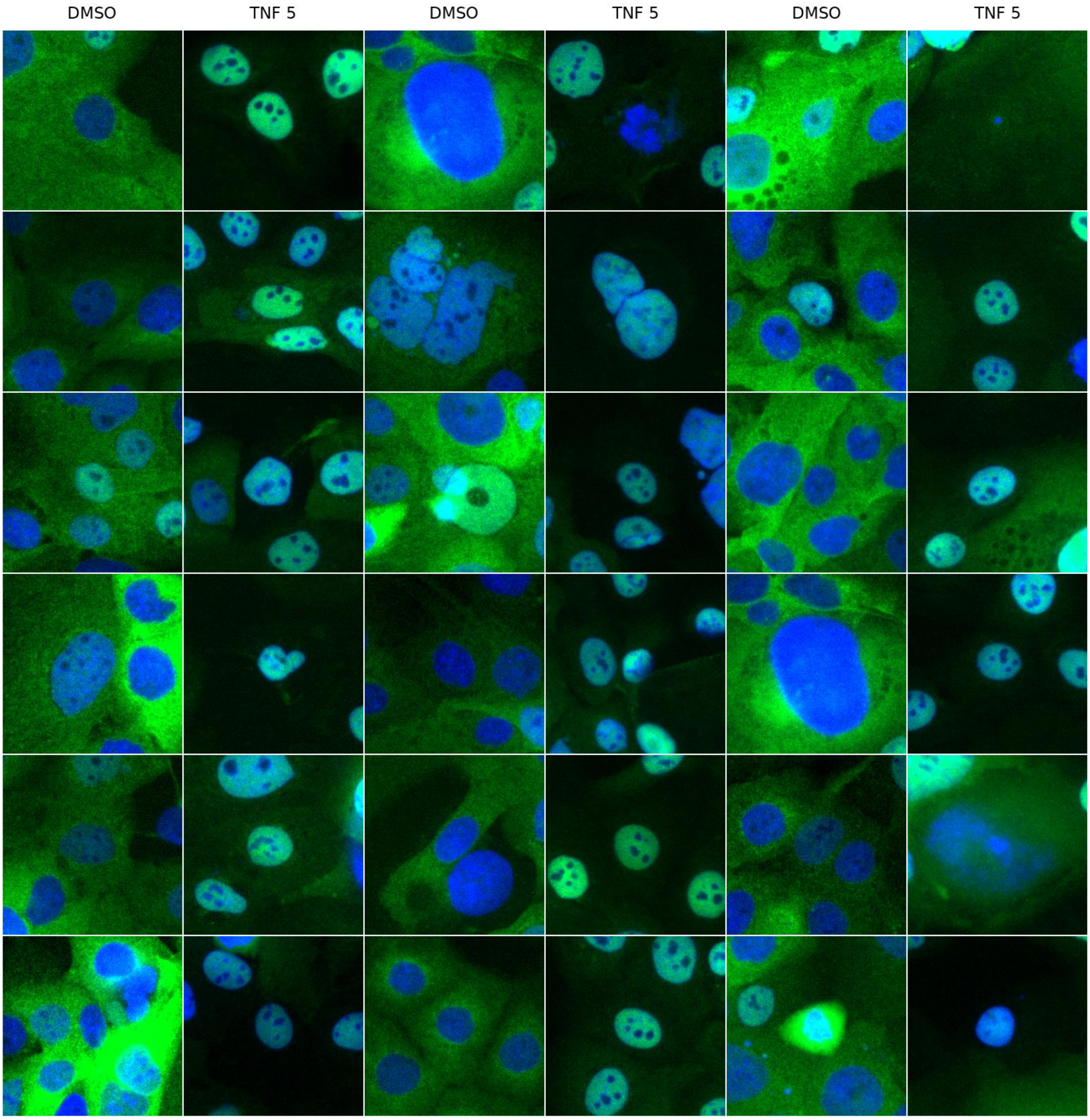


**Supplementary figure 3** Random samples of real cropped images of cells from the nuclear translocation assay. Concentration is 5 μM.

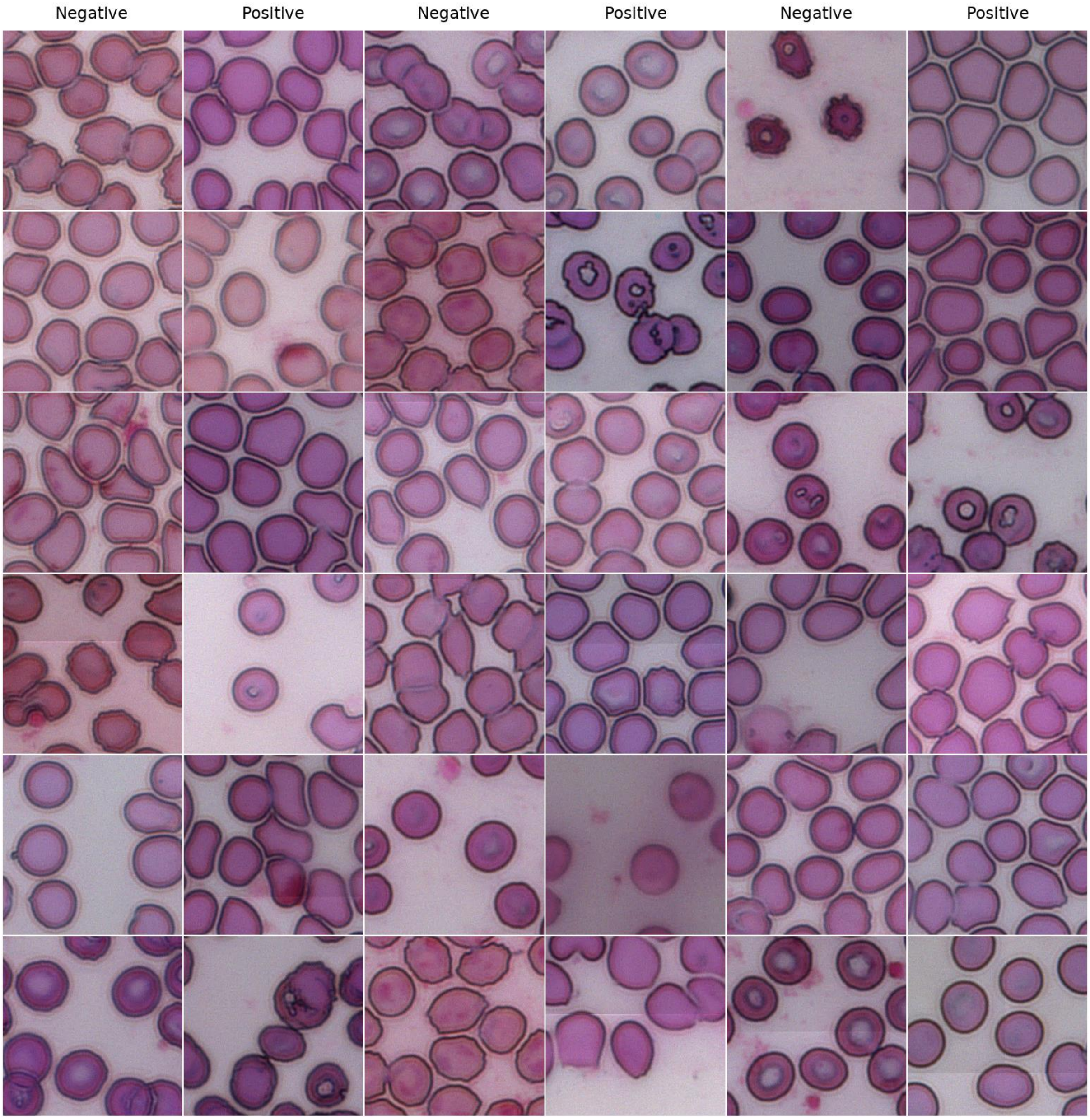


**Supplementary figure 4** Random samples of real cropped images of cells from thin blood smears, negative or positive to a qPCR test against Malaria, but assessed as negative by a microscopist, that is without visible parasites.

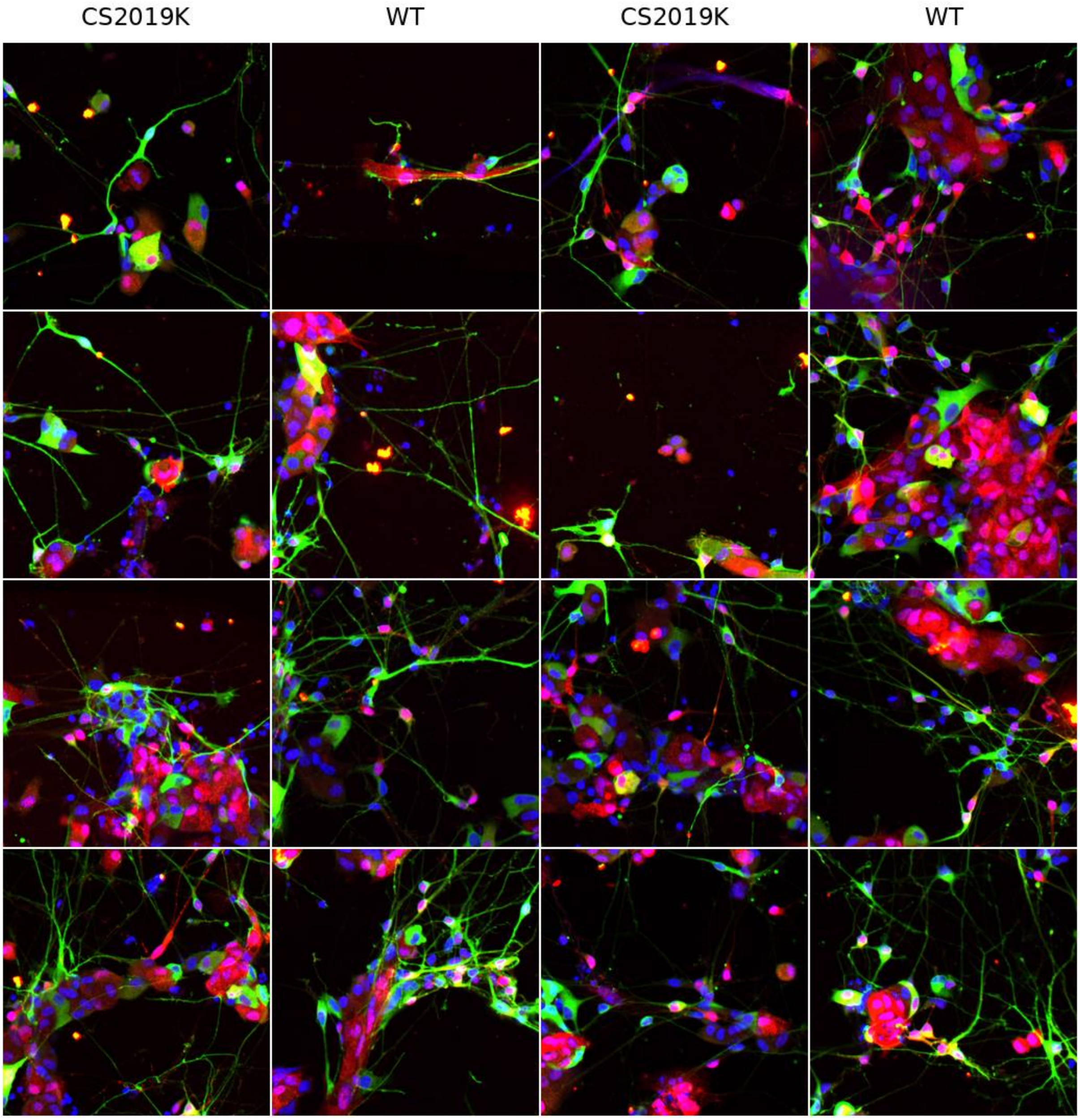


**Supplementary figure 5** Random samples of real cropped images from the Parkinson-LRRK2-CS2019K assay

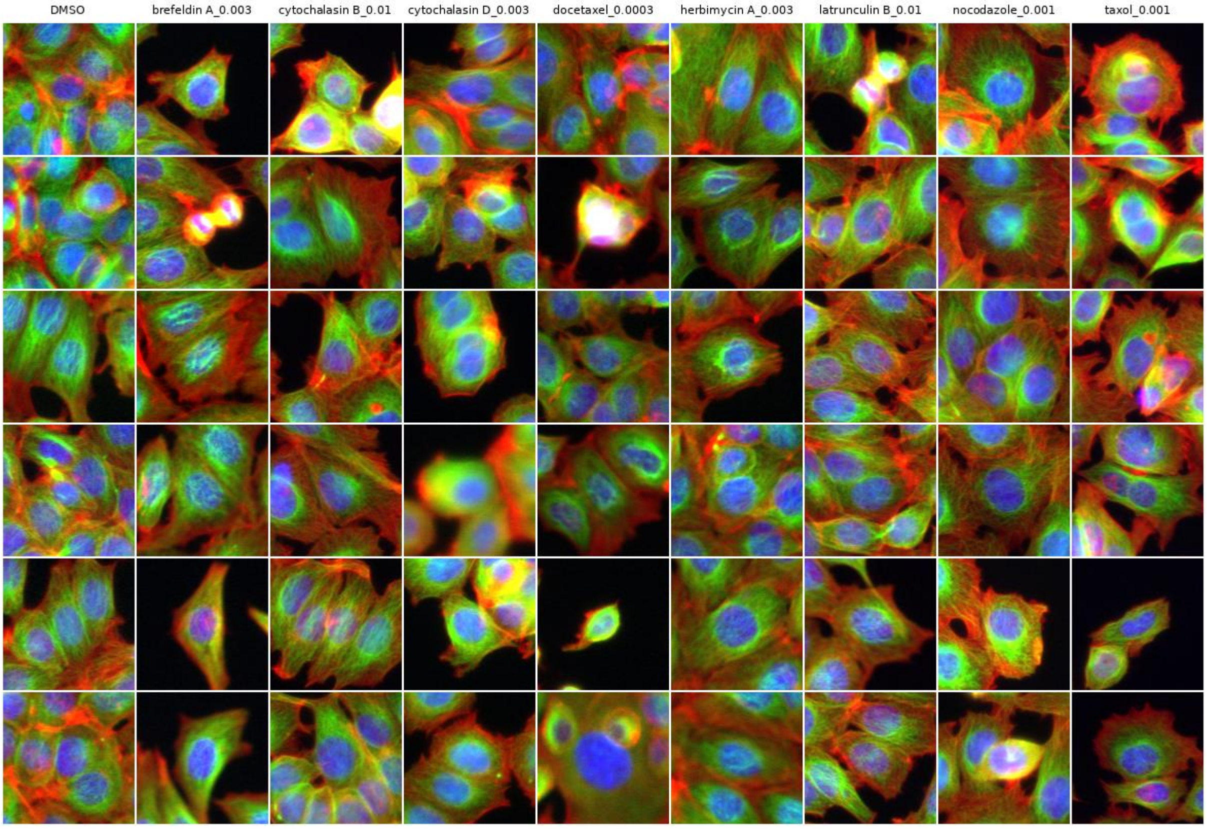


**Supplementary figure 6** Random samples of real cropped images of cells from low concentration compound treatments from the BBBC021 dataset (concentration unit is μM).

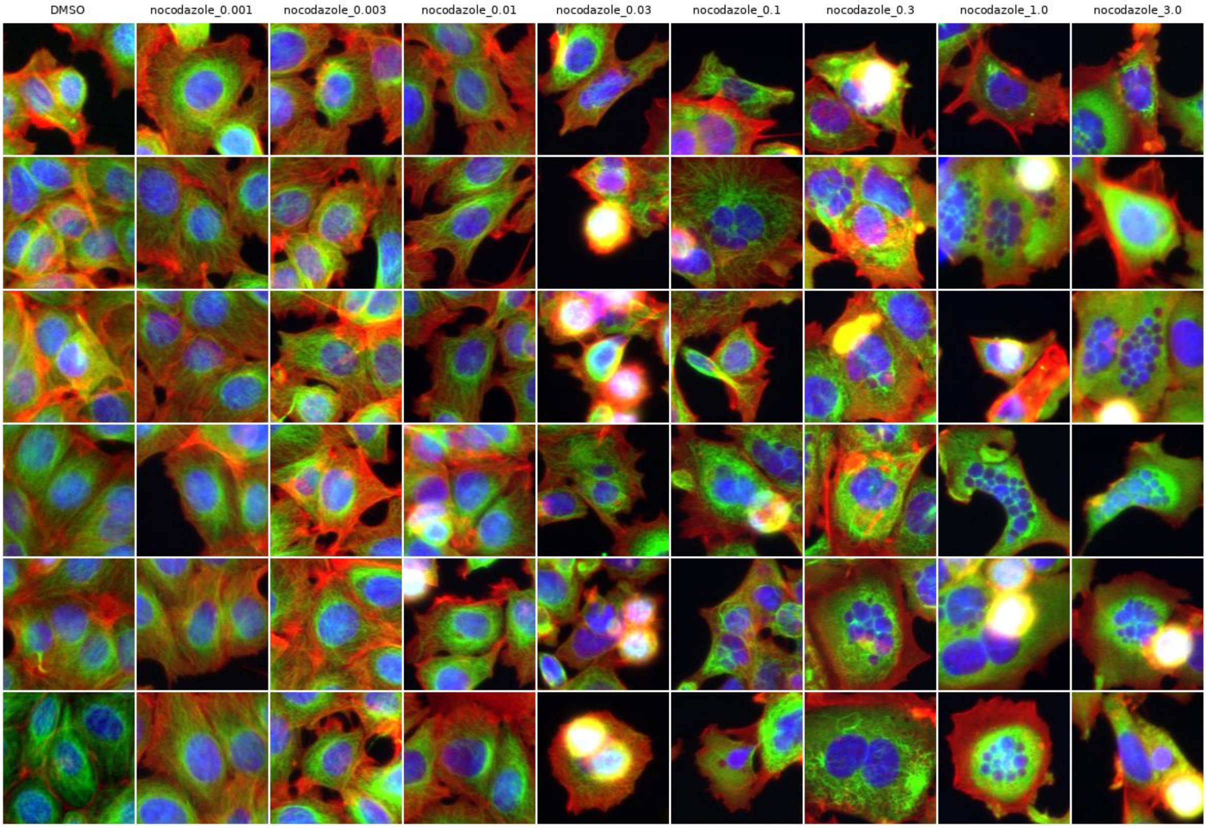


**Supplementary figure 7** Random samples of real cropped images of cells from each concentration of a dose response treatment of Nocodazole from the BBBC021 dataset (concentration unit is μM).

| **Dataset** | **Concentration (µM)** | **Mean accuracy** | **SD accuracy** |
|---|---|---|---|
| DMSO vs Brefeldin A | 0.003 | 0.83 | 0.02 |
| DMSO vs Brefeldin A | 10.0 | 0.97 | 0.03 |
| DMSO vs Cytochalasin B | 0.01 | 0.83 | 0.08 |
| DMSO vs Cytochalasin B | 30.0 | 0.94 | 0.01 |
| DMSO vs Cytochalasin D | 0.003 | 0.91 | 0.09 |
| DMSO vs Cytochalasin D | 10.0 | 0.97 | 0.02 |
| DMSO vs Docetaxel | 0.0003 | 0.83 | 0.03 |
| DMSO vs Docetaxel | 1.0 | 0.88 | 0.05 |
| DMSO vs Herbimycin A | 0.003 | 0.81 | 0.05 |
| DMSO vs Herbimycin A | 10.0 | 0.86 | 0.04 |
| DMSO vs Latrunculin B | 0.01 | 0.78 | 0.02 |
| DMSO vs Latrunculin B | 30.0 | 0.96 | 0.01 |
| DMSO vs Nocodazole | 0.001 | 0.82 | 0.04 |
| DMSO vs Nocodazole | 3.0 | 0.93 | 0.02 |
| DMSO vs Taxol | 0.001 | 0.86 | 0.00 |
| DMSO vs Taxol | 3.0 | 0.93 | 0.02 |
| Malaria (qPCR- vs qPCR+) | n/a | 0.74 | 0.03 |
| LRRK2 (G2019S vs WT) | n/a | 0.63 | 0.03 |

**Supplementary figure 8** Accuracy of a basic CNN classifier on the datasets used in this paper. For each dataset, we trained a classifier with 4 convolutional layers for 10 epochs using a 4-fold cross-validation, and computed the mean and standard deviation of the accuracy over these 4 runs. For the BBBC021 datasets, training was performed for the lowest and highest concentrations of each compound treatment. Even in the latter case, where differences could not be assessed by eye, good accuracy could still be reached quickly, indicating that CNN can leverage invisible discriminative features. For the LRRK2 dataset, each well of a well plate was split in many tiles with the corresponding well annotation to feed the CNN. For the Malaria dataset, random tiles from each thin blood smear were used to feed the CNN with the corresponding thin blood smear annotation.

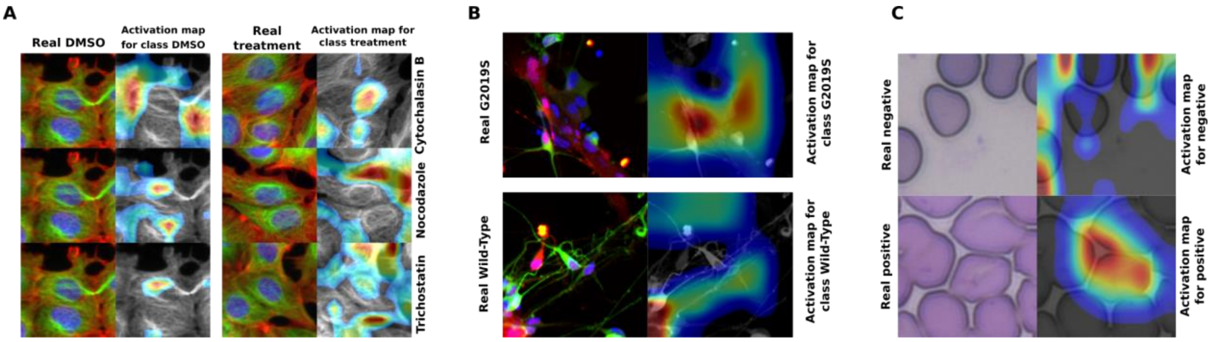


**Supplementary figure 9** Class Activation Maps for classifiers trained on 3 of our datasets. Activation Maps show which areas were considered important by the network, but they do not show precisely what specific difference of signal triggered discrimination, which, in case of subtle phenotype, makes interpretation impossible.
